# cDNA display coupled with next-generation sequencing for rapid activity-based screening: Comprehensive analysis of transglutaminase substrate preference

**DOI:** 10.1101/2021.09.08.459404

**Authors:** Jasmina Damnjanović, Nana Odake, Jicheng Fan, Beixi Jia, Takaaki Kojima, Naoto Nemoto, Kiyotaka Hitomi, Hideo Nakano

**Author notes:** To whom correspondence should be addressed: Jasmina Damnjanović, Graduate School of Bioagricultural Sciences, Nagoya University, Furo-cho, Chikusa-ku, Nagoya 464-8601, Japan.

## Abstract

cDNA display is an *in vitro* display technology based on a covalent linkage between a protein and its corresponding mRNA/cDNA, where a stable complex is formed suitable for a wide range of selection conditions. A great advantage of cDNA display is the ability to handle enormous library size (10^12^) in a microtube scale, in a matter of days. To harness its benefits, we aimed at developing a platform which combines the advantages of cDNA display with high-throughput and accuracy of next-generation sequencing (NGS) for the selection of preferred substrate peptides of transglutaminase 2 (TG2), a protein cross-linking enzyme. After the optimization of the platform by the repeated screening of binary model libraries consisting of the substrate and non-substrate peptides at different ratios, screening and selection of combinatorial peptide library randomized at positions -1, +1, +2, and +3 from the glutamine residue was carried out. Enriched cDNA complexes were analyzed by NGS and bioinformatics, revealing the comprehensive amino acid preference of the TG2 at targeted positions of the peptide backbone. This is the first report on the cDNA display/NGS screening system to yield comprehensive data on TG substrate preference. Although some issues remain to be solved, this platform can be applied to the selection of other TGs and easily adjusted for the selection of other peptide substrates and even larger biomolecules.

## Introduction

Transglutaminases (TGs: EC 2.3.2.13) are enzymes catalyzing transamidation, a transfer reaction between an acyl donor (peptidyl glutamine) and an acyl acceptor (amino group of lysine). As a result, a stable isopeptide bond is formed, resistant to proteolytic degradation. Owing to their transamidation activity, TGs are known for their role in cross-linking of proteins and peptides. Since transamidation proceeds *via* the formation of acyl-enzyme intermediate, which is a rate-limiting step, TGs show high specificity towards acyl donors, while reacting on a variety of acyl acceptors such as lysines of proteins and peptides and even low molecular weight amines.^(5)^

The presence of TGs is vast across kingdoms of plants, animals and microorganisms, with many of them having been studied and characterized. In mammals, eight different types of TGs have been identified (TG1, TG2, TG3, TG4, TG5, TG6, TG7, and factor XIII), with functions ranging from blood clotting, epidermis and hair follicle formation, wound healing, apoptosis, extracellular matrix formation and cell adhesion.^(8)^ Among mammalian TGs, transglutaminase 2 (TG2) is the most-studied and yet most elusive TG because of its omnipresence in cellular compartments and tissues and its multifunctionality which includes transamidase, GTPase, protein disulfide bond isomerase and protein kinase activity.^(9; 30)^ All mammalian TGs are homologous and require calcium ion for activation. Aberrant expression and function of TGs have serious effects on human health and cause conditions such as hemorrhage, celiac disease, cancer, fibrosis, Alzheimer’s and Huntington’s disease, and lamellar ichthyosis.^(18)^ Although extensively studied, more work is needed to fully understand the biological function of TGs for which sensitive and specific *in situ* detection of TG activity is desired. Artificial fluorescently labeled glutamine (Gln)-peptide probes (‘Hitomi peptides’), have greatly contributed to *in vitro* and *in situ* detection and measurement of TG activity.^(29; 28; 33; 7; 21; 19)^ These probes have been developed by the selection from random peptide libraries by phage display technology, and are now available for studies of TG isozymes. Up to date, these probes have not been optimized in terms of amino acid preference at Gln-surrounding positions or used to investigate TG substrate preference by protein engineering.

Protein engineering is an indispensable set of tools developed to tailor the protein and peptide function for specific needs. However, screening of large libraries has long been a bottleneck in terms of time, labor and cost. For example, random mutagenesis of five amino acid residues generates a diversity of 10^6^ at an amino acid level and 10^9^ at a nucleotide level. With six amino acid residues, the diversity increases to 10^7^-10^10^. Having the numbers in mind, it is easy to understand why an efficient screening system is necessary.

*In vitro* screening and selection methods utilize cell-free protein synthesis and various display technologies to physically link genotype and phenotype. Compared to *in vivo* methods where library size is limited by the transformation of DNA into *E. coli* or yeast, *in vitro* methods do not require transformation and thus allow for larger library size and, selection of cytotoxic proteins with less processing time. Widely used *in vitro* methods include mRNA/cDNA display, ribosome display and phage display, although phage display includes an *in vivo* step. mRNA/cDNA display was pioneered by two groups at a similar time, Roberts and Szostak’s group^(24)^ and Nemoto and Yanagawa’s group^(23)^. This technology relies on the formation of a covalent link between the genotype, represented by mRNA/cDNA and phenotype, represented by the corresponding protein, via puromycin. The convenience of complete control of expression/screening conditions, stable genotype-phenotype linkage, incorporation of unnatural amino acids and handling of large libraries makes mRNA/cDNA display a preferred method for screening and selection in protein engineering.

Numerous reports exist on mRNA/cDNA display being applied to affinity-based screening and selection of peptides and antibody fragments.^(2; 1; 17; 15; 22; 31)^ Utilization of mRNA/cDNA display for activity-based selection of enzyme substrates has been described, however to a much less extent. The representative applications include analysis of substrate scope of protein-modifying enzymes in proteomics research, such as caspase using the library of the mRNA-displayed human proteome ^(12)^, viral protease and a kinase ^(14)^, and metalloprotease ^(27)^. Very recently, this technology has been applied to study the substrate scope of enzymes involved in the modification of post-translationally modified peptides ^(6; 32)^ demonstrating that the substrate preference can be studied in great detail by mRNA/cDNA display. These achievements clearly show the great potential of mRNA/cDNA display for studies of other biologically and industrially relevant enzymes and their substrates. Aiming to study the substrate specificity of microbial transglutaminase (MTG) and design its artificial substrate for biotechnological applications, Lee *et al*. used mRNA display to isolate a novel Gln substrate of MTG after six rounds of screening and selection from a 10-mer random library ^(16)^.

To expand the potential of mRNA display in TG-related research, we aimed to develop a cDNA display/NGS platform for comprehensive analysis of the TG substrate preference (Fig. 1) which would encompass the majority of the reactive sequences rather than the best hits, enable their ranking and analysis of the consensus sequence. We also aimed to reduce the time and labor associated with multiple selection rounds and probe the sensitivity of the platform in a single selection round. For the platform development, we used TG2 and its substrate, T26 peptide, previously isolated from a phage-displayed 12-mer-peptide library.^(29)^ We first optimized *in vitro* synthesis and display of the T26 peptide and selection conditions by the repeated screening of binary model libraries consisting of T26 and non-substrate peptide T26A (Gln is replaced with Ala). In the following, our platform was deployed for screening of a T26-based random library, LibQ, with Gln randomized by NNK codon, to confirm the efficiency of the activity-based screening. In continuation, a T26-based library, Lib4, with residues at positions -1, +1, +2 and +3 from Gln randomized by NNK codon was constructed and screened by our platform. By analyzing the enriched sequence data, we obtained a consensus sequence indicating the TG2 preference at randomized positions of the peptide backbone. This result demonstrates that our platform represents a time-, labor- and cost-efficient tool for comprehensive analysis of TG substrate preference and development of TG peptide probes.

**Figure 1.**
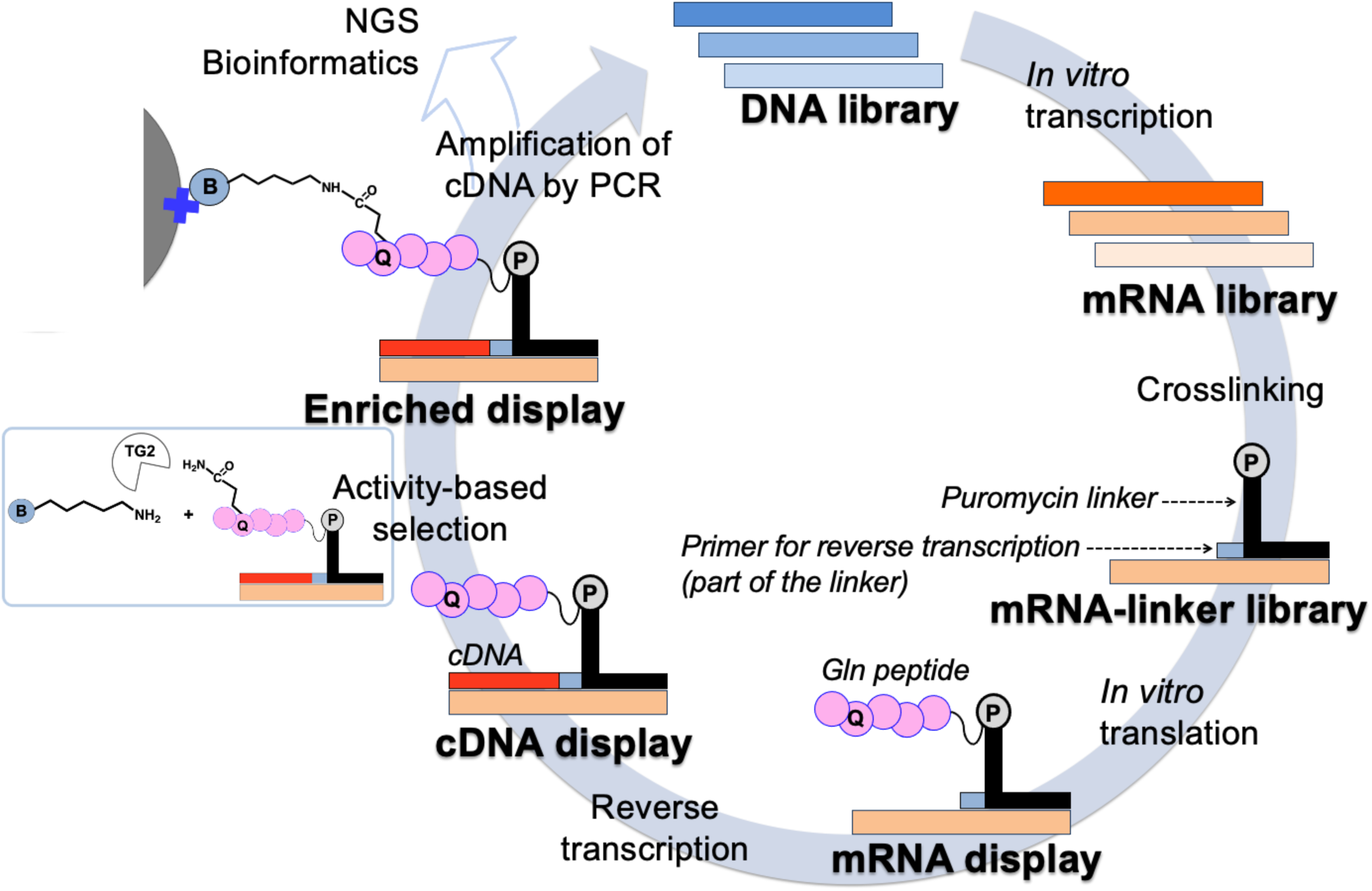
Outline of the cDNA display platform for screening and selection of preferred TG peptide substrate. During the selection step, glutamine of the reactive displayed peptides is biotinylated in TG-catalyzed crosslinking reaction between the amino group of the pentylamine-biotin and side-chain of glutamine (as indicated by the small squared image and illustration of the enriched display). Biotinylated display complexes are collected by the streptavidin-coated magnetic beads (colored gray).

## Results

### T26 peptide display

We first attempted to display T26 peptide with a C-terminal GST-tag (Fig. S1), since this construct ensured soluble production of T26 in *E*. *coli* cells previously ^(29)^. However, the selection of T26-GST display from binary model libraries beyond 1:1 molar ratio of T26-GST and negative control peptide was unsuccessful (Fig. S2). The suspected reason is the aggregation propensity of the GST tag under the conditions used for mRNA display generation and manipulation (data not shown).

We next tested the display of the untagged T26 peptide (Fig. S1, Table S1). mRNA display (peptide-linker-mRNA complex) of untagged T26 was produced by the PURE*frex* system and checked for solubility. Surprisingly, the display was fully soluble and the efficiency of mRNA display synthesis proceeded with approx. 70% yield (Fig. 2), as judged by the band density of mRNA-linker complex and mRNA display complex. The untagged T26 construct (abbreviated as T26) was used in consequent experiments and for the construction of peptide libraries.

**Figure 2.**
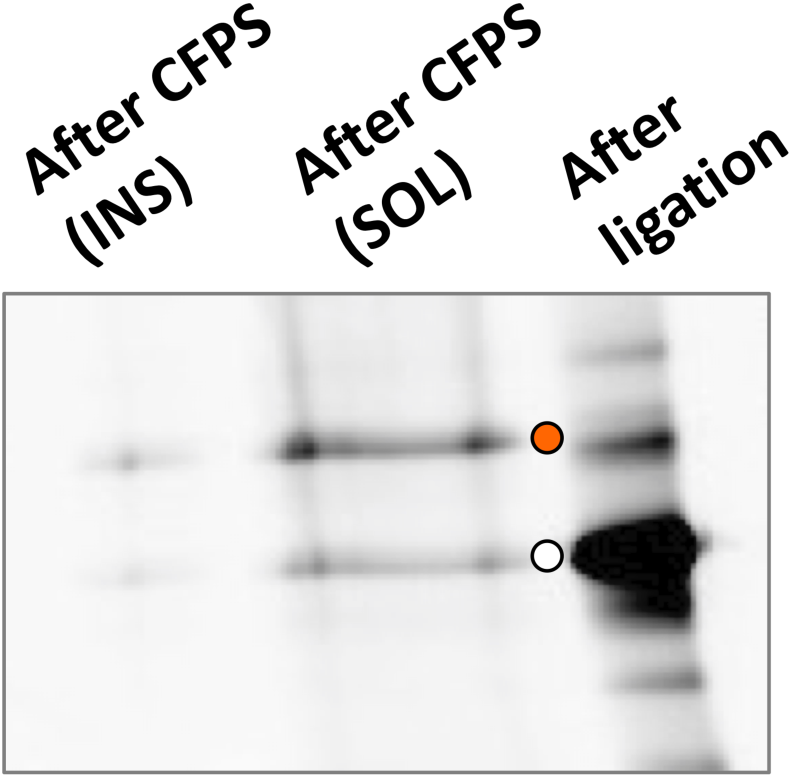
Urea SDS-PAGE gel of the ligation product and mRNA display complex of the untagged T26 construct visualized using fluorescence imager with FITC filter. Orange circles indicate the position of mRNA display complex and white circles indicate the position of the ligation product.

### Generation of binary model libraries

Binary model libraries containing T26: T26A (non-substrate peptide) = 1:4 or 1:50 were made by mixing corresponding DNA in specified molar ratios and were used as templates for mRNA library generation. mRNA synthesis in 30-μL scale proceeded with the overall yield of 29 μg for 1:4 mRNA library and 55 μg for 1:50 mRNA library. Hybridization and photo-crosslinking yielded mRNA-linker complexes of both libraries (Fig. 3A). These complexes were used as templates for cell-free protein synthesis and yielded mRNA display complexes (Fig. 3B).

**Figure 3.**
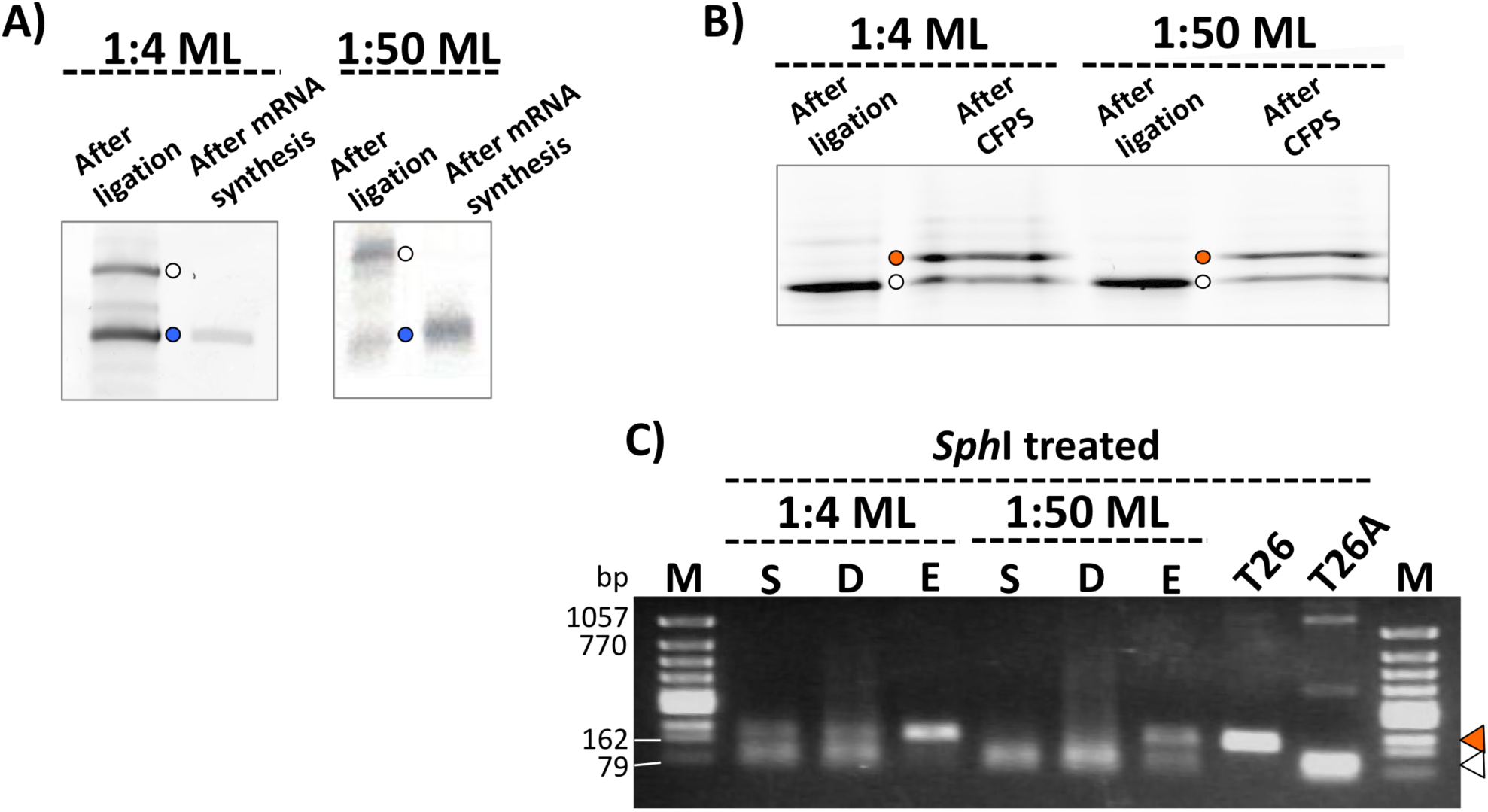
Preparation of binary model libraries and selection of T26. (A) Urea PAGE of generated mRNA libraries and corresponding ligation products detected with fluorescence imager after staining with SYBRGold. Blue circles indicate the position of mRNA and white circles indicate the position of ligation products. (B) Urea SDS-PAGE of ligation products and mRNA display complexes detected by fluorescence imager under the FITC filter. Orange circles indicate the position of mRNA display complexes and white circles indicate the position of ligation products. (C) Electrophoresis of amplified and *Sph* I-treated DNA samples of the original (D), selected (E) and leftover (S) library. Orange triangles indicate the position of the T26 DNA band and white triangles indicate the position of the T26A DNA band.

### Selection of T26 sequence from binary model libraries

After the selection and subsequent processing to obtain enriched DNA on the surface of magnetic beads, cDNA of the original, enriched and leftover library was PCR-amplified by step 1 of the nested PCR, digested with *Sph* I and analyzed by agarose gel electrophoresis, (Fig. 3C). Band density analysis indicates enrichment factors of 5 and 30 for 1:4 and 1:50 libraries respectively.

### Generation of random libraries

The display of random libraries, LibQ (Gln position randomized by NNK codon) and Lib4 (positions -1,+1,+2,+3 from Gln randomized by NNK codon), started with the preparation of the DNA libraries from chemically synthesized ssDNA and their conversion into the corresponding mRNA libraries. mRNA synthesis at 30-μL scale proceeded with the overall yield of 36 μg for Lib4 mRNA library and 56 μg for LibQ mRNA library. The formation of mRNA-linker complexes was confirmed for both libraries (Fig. 4A). Obtained mRNA-linker complexes were used for cell-free protein synthesis as templates and yielded mRNA display complexes (Fig. 4B).

**Figure 4.**
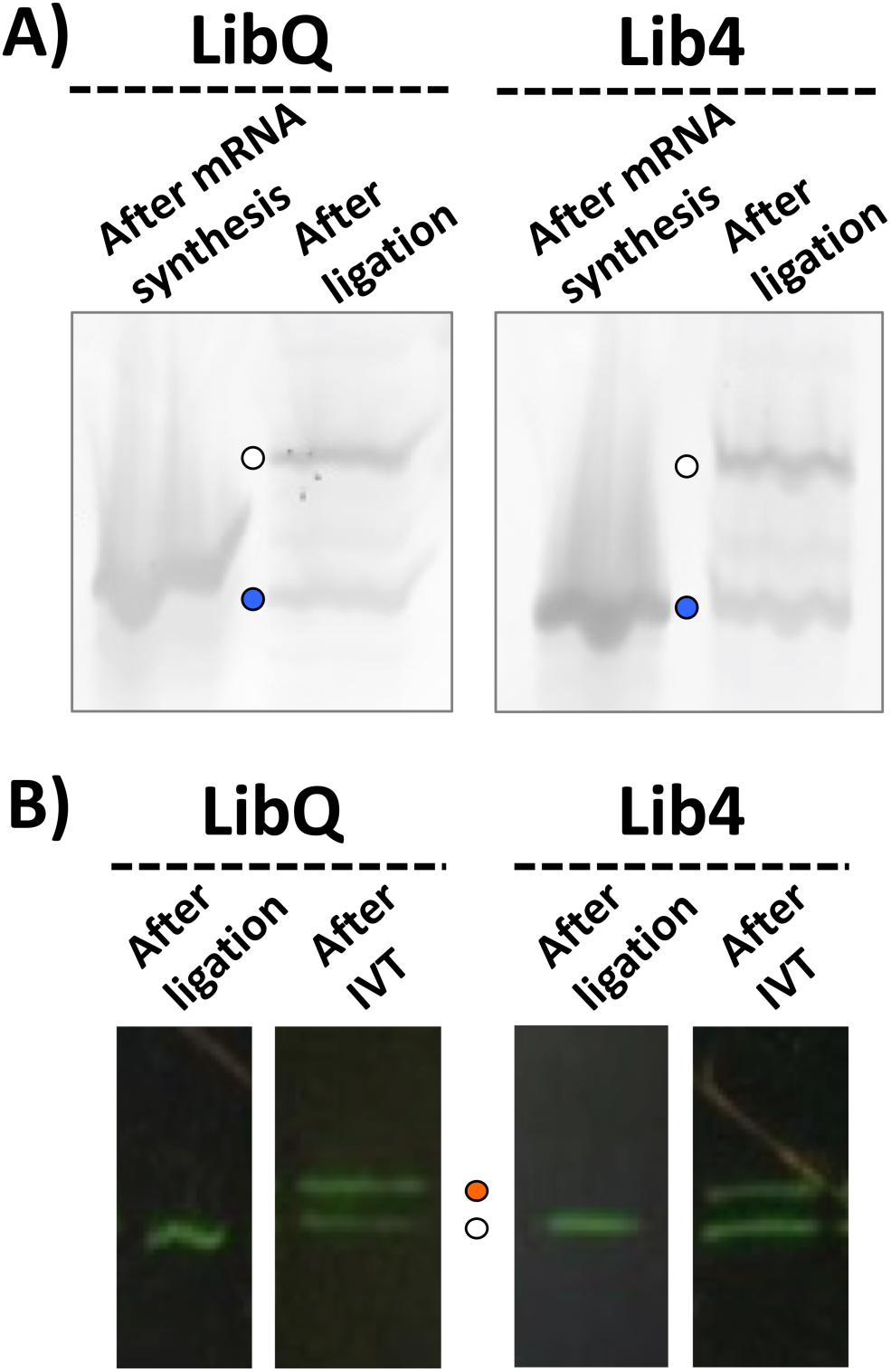
Preparation of random libraries. (A) Urea PAGE of generated mRNA libraries and corresponding ligation products detected with fluorescence imager after staining with SYBRGold. Blue circles indicate the position of mRNA and white circles indicate the position of ligation products. (B) Urea SDS-PAGE of ligation products and mRNA display complexes detected by fluorescence imager under the FITC filter. Orange circles indicate the position of mRNA display complexes and white circles indicate the position of ligation products.

### Selection of peptide sequences from random libraries

The selection and subsequent processing to obtain enriched DNA on the surface of magnetic beads proceeded in the same way as for the binary model libraries. cDNA of the original, enriched and leftover library was PCR-amplified by step 1 of the nested PCR, and analyzed by agarose gel electrophoresis. Bands corresponding to the original and leftover libraries were observed (Fig. 5A). Since the corresponding bands were not observed for the enriched library, step 2 of the nested PCR was performed with different volumes of the PCR reaction mixture from step 1 as a template. After step 2, bands corresponding to the DNA of the enriched library were detected (Fig. 5B). We believe that due to the low amount of the reactive peptide sequences, an initial amount of cDNA enriched on the beads was too low to obtain a detectable amplification product after only one round of PCR. The enriched DNA was purified, modified for sequencing and subsequent read processing, and analyzed by NGS.

**Figure 5.**
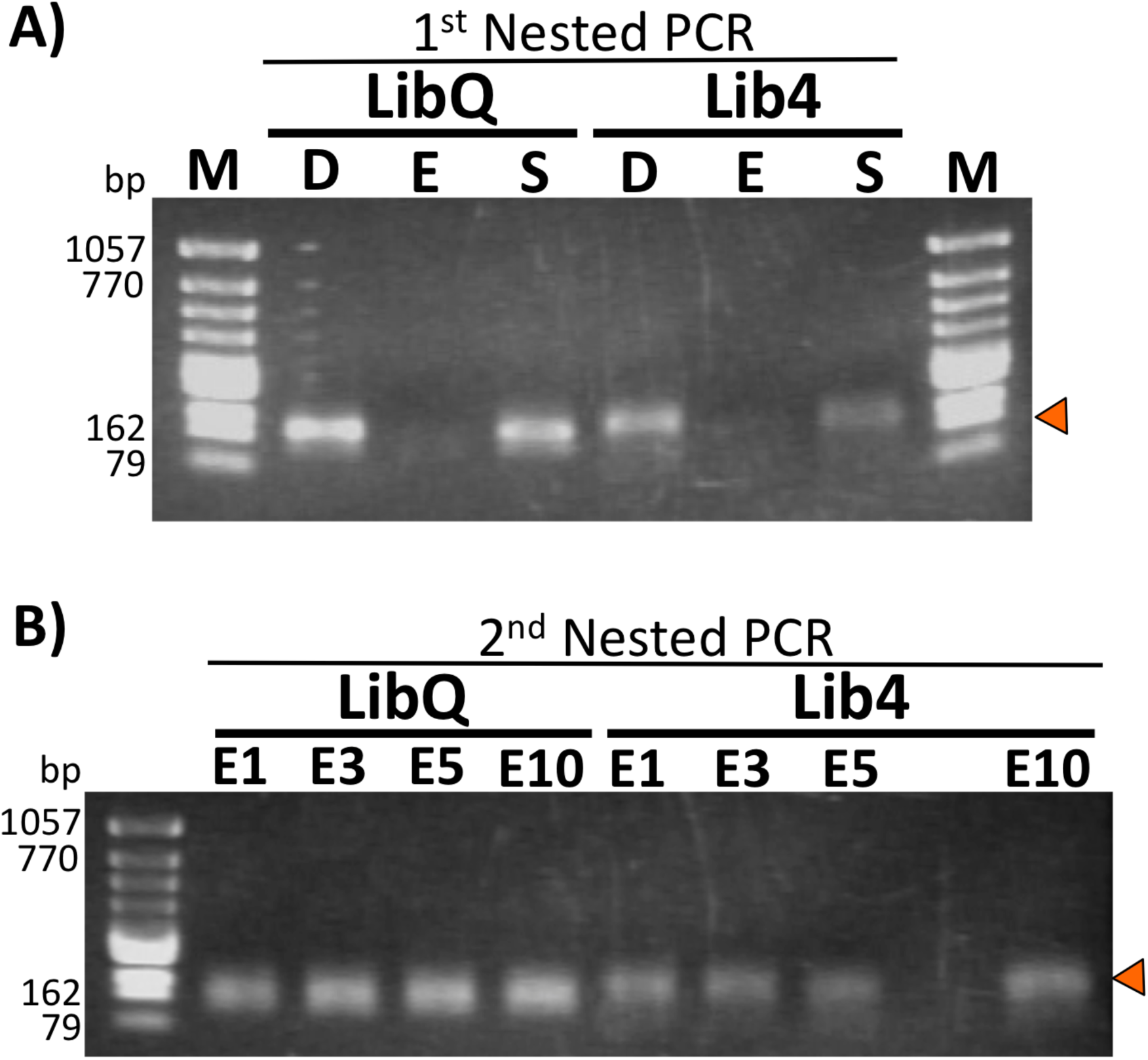
(A) Electrophoresis after PCR amplification (first step of the nested PCR) of DNA from the original (D), selected (E) and leftover (S) LibQ and Lib4 libraries. (B) Electrophoresis after PCR amplification (second step of the nested PCR) of the selected (E) LibQ and Lib4 libraries. Lanes E1, E3, E5 and E10 show the bands of the PCR products after the second step of the nested PCR when 1, 3, 5, and 10 μL of the first PCR reaction was used as a template respectively. The orange triangle indicates the position of amplified DNA bands.

### Analysis of the sequencing data from random libraries

The raw NGS data were analyzed as described in Materials and Methods. After converting the processed DNA sequence data into the amino acid sequence data, the total number of sequences in both libraries, before and after selection, was calculated along with the number of identical sequences at each mutated amino acid position. Based on these numbers, the enrichment factor at each position was calculated. The enrichment factor of Gln from LibQ was approx. 3 (Fig. 6). The enrichment factor of the original T26 sequence from Lib4 was approx. 2. However, in Lib4, residues other than the original ones have also been enriched with similar enrichment factors. The data (Fig. 7) indicate that His and Gln were predominantly enriched in position -1, Ser and Cys at position +1, Tyr at position +2, and Val and Ile at position +3. Based on the enrichment data, the consensus sequence was defined as H/Q-Q-S/C-Y-V/I-D-P-W-M-L-D-H. The list of top 100 sequences enriched from Lib4 and ranked by the enrichment factor is given in Table S2. Among the top 100 enriched peptides, approx. 30%, including the first six sequences, contain Gln-Gln motif in positions -1 and 0, in accordance with the consensus sequence. In a few sequences, Gln-Gln motif is present at 0 and +1 positions. Interestingly, selected sequences also show the presence of Cys, mostly at positions -1 and +1, around the reactive Gln. In the top 100 sequences, position +1 is mostly occupied by Tyr (23/100) followed by Cys, Val and Phe (12/100 for Cys and Val and 14/100 for Phe). Position +2 shows less preference for any particular residue, while at position +3, Val and Ile are dominant, as also observed in the consensus sequence.

**Figure 6.**
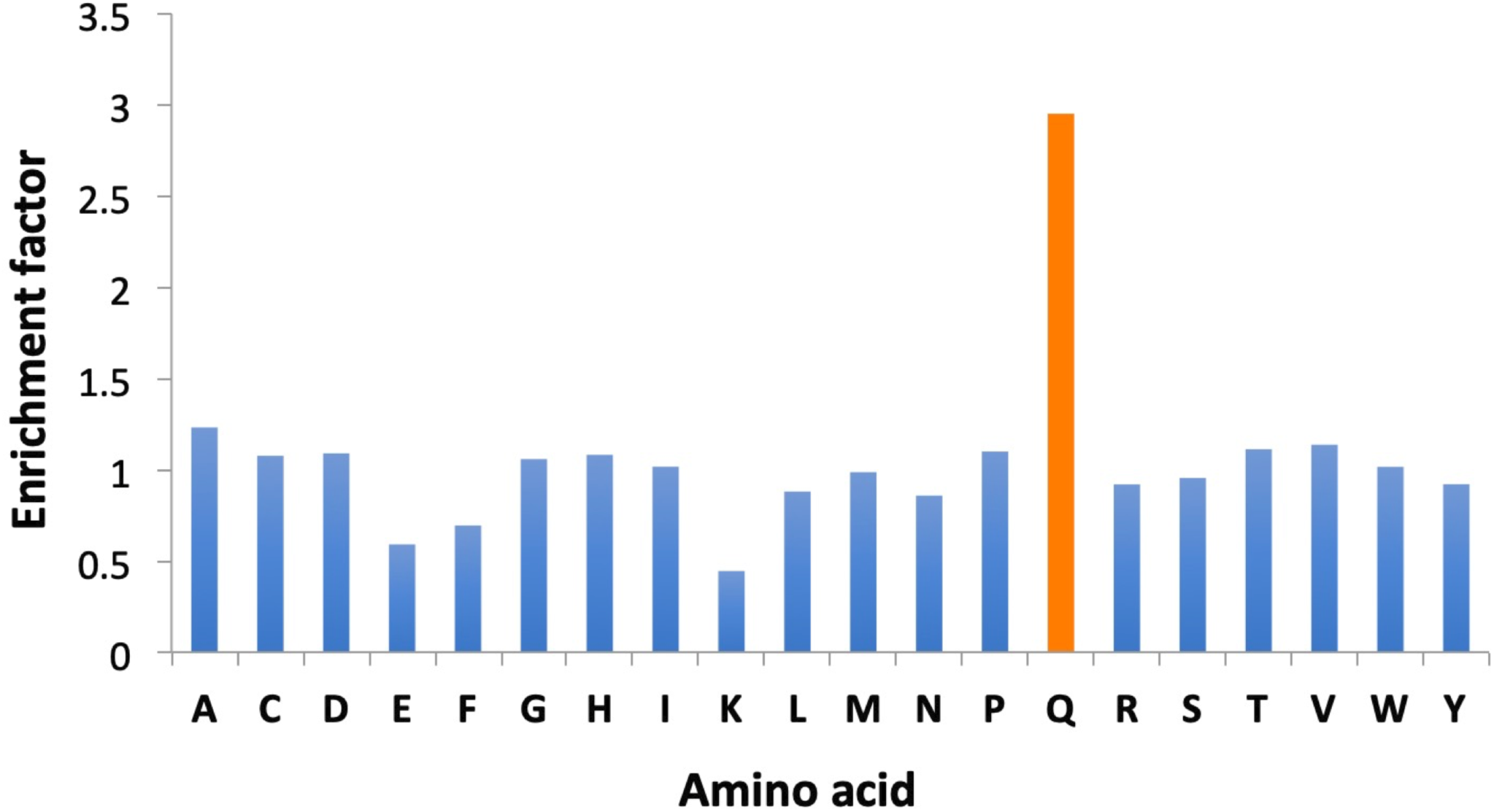
The enrichment factor of each amino acid residue at the randomized Gln position of T26 in LibQ.

**Figure 7.**
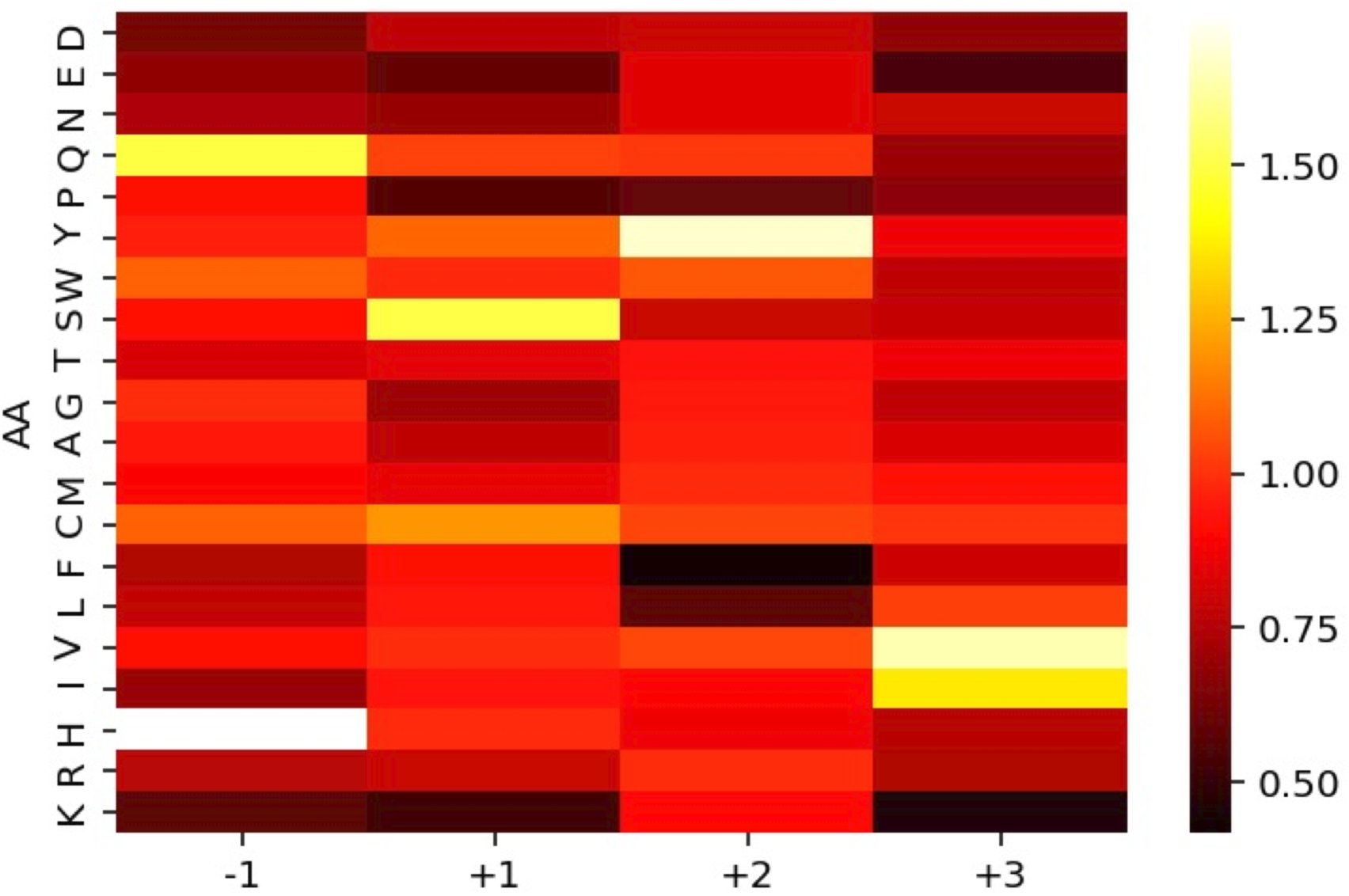
The heatmap showing the color-coded enrichment factor of each amino acid residue at four randomized positions (-1, +1, +2, +3) of T26 in Lib4.

### Identification of the new potential protein targets of TG2

We used NCBI Blastp (https://blast.ncbi.nlm.nih.gov/Blast.cgi) search with the first seven residues of the enriched peptide sequences (T26, or combination of other enriched residues, QQCYIDP) as a query against a non-redundant sequence database of mouse and human proteins to verify if potential new protein targets of TG2 can be identified based on the presence of the query sequence motifs in their primary structure. Table 1 summarizes the search results. Both sequences have matching motifs in proteins with diverse functions. The localization of these potential targets also matches the one of TG2. Interestingly, both searches converge towards variable and junction regions of immunoglobulins, indicating a possible role of TG2 in the regulation of antibody function. To the best of our knowledge, this role of TG2 has not been scientifically described yet.

**Table 1.** Summarized results of the Blastp search with two query sequences, consisting of the first 7 a.a. residues of the enriched peptides against the human and mouse proteins. Only the matches with 6-7 residues including one residue mismatch, or 5 residues without a mismatch are given. If aligned amino acid residues are not identical but are similar in size and nature, this was not considered a mismatch.

| Query | T26 (HQSYVDP) | QQCYIDP |
| --- | --- | --- |
| Match | Ig light chain junction region<br>Ig heavy chain junction region<br>Ig kappa light chain variable region<br>Ankyrin-3 isoforms<br>mCG144990 | Zinc transporters (solute carrier 30 family)<br>Ig kappa light chain variable region<br>Ig light chain junction region<br>Heparanase 2 and 3<br>Coatomer subunit beta<br>Telomere length regulation protein TEL2 homologs<br>Gamma-glutamyl hydrolase precursor<br>mCG12390 |

## Discussion

The platform for rapid screening and comprehensive analysis of TG substrate preference has been established and used for elucidation of the substrate profile of TG2. The platform relies on in *vitro* transcription and translation, which saves time and increases workable library size while reducing the risk of positive hit loss due to transformation efficiency or low expression level. For mRNA/cDNA display generation, we used the PURE system^(26)^, which contains only necessary components of the transcription/translation machinery in a highly pure state, thus enabling the tight control of the expression conditions and eliminating the risk of proteolytic degradation. To facilitate the workflow, we used a puromycin linker for mRNA-protein chemical crosslinking developed by the Nemoto group^(20)^ to ensure the fast and stable formation of the mRNA/cDNA display complexes with minimum risk to loss of genetic information during experimental processing.

We combined cDNA display technology with NGS to yield a system for the rapid activity-based evolution of proteins and peptides from large libraries in the shortest time with minimal demand for labor and, with full control over the expression conditions. The use of NGS increases the throughput and enables information-driven protein evolution since we can monitor the sequence enrichment after each screening cycle and gather broad information on sequence evolution.

Having in mind the great importance and abundance of TGs, we have chosen to develop a system to study their substrate profile as our first goal. A previous study reported about the preferred and specific substrate of TG2, T26 peptide, isolated from a random peptide library screened by phage display.^(29)^ Follow-up study revealed that positions +2, +3, +4 play an important role in the reactivity of T26^(10)^, however comprehensive substrate profiling of TG2 has not been performed yet. T26-derived peptide libraries were displayed in untagged form with high translation efficiency and complete solubility of mRNA/cDNA display complex. It is believed that the formation of the mRNA display complex itself helps the peptide stay soluble and available for the enzymatic reaction. The binary model libraries consisting of T26 and T26A peptides have been used to optimize the selection and post-selection treatment of the beads with an enriched cDNA display library. Our first enrichment results indicated that non-specific interactions between display complexes and the surface of magnetic beads during selection cause serious problems in the quality of enriched DNA. This was manifested as the presence of T26A DNA in the enriched sample alongside T26 DNA (data not shown). To solve the problem, we used several strategies among which extensive washing with the buffer containing a high concentration of salt and detergent proved critical. In the end, after a single round of selection, we achieved five times enrichment of the T26 sequence from 1:4 and 30 times enrichment from the 1:50 model library. This result demonstrates the applicability of our platform for screening and selection of TG substrate preference, and we, therefore, proceeded to screen random libraries under the same conditions.

Analysis of enriched DNA from LibQ indicated that Gln was predominantly enriched, as expected. Lower-than-expected enrichment factor for Gln is believed to be the consequence of performing a single selection round, and persistence of non-specifically enriched peptide sequences. However, since Gln was the only enriched sequence, with all other sequences having enrichment factors around 1 or below, we did not consider this an issue. As for the Lib4, the sequence corresponding to the residues of T26 peptide was enriched at all positions, in addition to the new alternative sequences at positions -1, +1, and +3. At position -1, the T26 residue, His, was enriched alongside Gln. Although the two residues are of a different chemical nature, other studies have also found the Gln-Gln and Gln-Gln-Gln motif in the preferred sequences of TG2^(13)^, as well as microbial TG^(16)^. These motifs are also present in natural protein targets of TG2, such as substance P, crystallin, and fibronectin. Asp, Glu, and Lys were the least preferred residues at position -1. A negative charge of Asp and Glu could lead to unfavorable interactions with the residues of the TG2 substrate-binding site. Lys could become acyl acceptor of the neighboring Gln, which would lead to self-cross-linking of the peptide. Thus, Lys is also expected to be among the least preferred residues at this position. At position +1, the T26 residue, Ser, was enriched alongside Cys. The two residues have a similar shape, size and chemical nature, thus it does not surprise that the two sequences share the similar preference of the enzyme. Interestingly, the presence of Cys in the peptide sequence did not prove harmful to the enzyme’s active site Cys, possibly due to the reducing environment of the selection reaction. This time as well, Lys was among the least preferred residues, likely for the same reason as in position -1. Besides Lys, among the least preferred residues were also Pro and Glu. At position +2, the T26 residue, Tyr, was preferably enriched. Meanwhile, the least preferred residues were Phe, Leu and Pro. This result implies that the hydroxyl group of Tyr is critical for the favorable enzyme-substrate interaction. At position +3, the T26 residue, Val, was enriched alongside Ile. The two enriched amino acids have a similar size and chemical nature, which indicates that small hydrophobic residues are preferred at this position. In contrast, the least preferred residues were charged Lys, Asp and Glu.

Besides the consensus sequence, we have also analyzed the top 100 enriched peptides ranked by the enrichment factor. Their sequences are well aligned with the consensus sequence however, show some differences. The top peptides show predominantly Gln at position -1 (28/100). The greatest difference between the consensus sequence and the sequences of the top 100 enriched peptides are the abundance of Tyr at position +1 (23/100 peptides), and the rather broad presence of hydrophobic and basic residues at position +2. This could be an indicator of the broader specificity of the enzyme at these two positions. Evidently, Tyr at position +1 or +2 is desired. Position +3 is dominated by Val and Ile, in the consensus sequence and in the sequence of top 100 enriched peptides. It should be noted that among the top 100 enriched peptides we have found seven sequences without Gln. Their presence in the original library could be explained by the errors in the chemical synthesis of ssDNA library fragments. Upon inspection of their sequences, we noticed that these peptides contain His and Trp near the N-terminus, which is a characteristic of streptavidin-binding peptides, meaning that they could bind to streptavidin and get falsely enriched during the selection. This phenomenon has been described before ^(29)^. In addition, some sequences in the top 100 list lack start codon, and these could only be explained by non-specific enrichment (no peptide expression but the mRNA-linker-cDNA complex gets bound to the streptavidin beads non-specifically).

Finally, the results of our Blastp search suggest that our platform can also be used for the identification of possible novel TG targets.

## Conclusion

The cDNA display/NGS platform for the selection of TG peptide substrates and analysis of TG substrate profile has been established. This *in vitro* system enables rapid screening and selection of preferred TG substrate peptides from random libraries based on enzymatic activity, resulting in enrichment of the selected sequences on the surface of the magnetic microbeads, from where they can be recovered and analyzed. Availability of the NGS can shorten the selection to only one or a few rounds, based on which we can deduce the substrate preference by proper data processing and analysis. We plan to use this system to screen for the substrate preference of other mammalian TGs and design appropriate peptide probes for detection and analysis of TG activity. Our hope is that the enriched sequence data can be also used to identify novel natural TG targets and promote TG-related research. The platform is further expected to become an indispensable tool for screening enzyme libraries during enzyme engineering.

## Materials and Methods

### Materials

Oligonucleotides were synthesized by Greiner Bio-One, Japan, or Integrated DNA Technologies, Singapore, or by Eurofins, Japan. Restriction enzymes, DNA polymerases (Pyrobest and PrimeStar), and Recombinant RNase Inhibitor were purchased from Takara Bio Inc., Japan. Recombinant Mouse Transglutaminase 2 was from Novus Biologicals, USA. Components of *in vitro* cell-free protein synthesis kit, PURE*frex*, were provided by GeneFrontier Corporation, Japan. Puromycin cnv-K and SBP linkers were obtained from Epsilon Molecular Engineering Inc., Japan.

### Preparation of T26 DNA constructs

Plasmids with the T26 gene constructs and their corresponding inactive (non-substrate) versions were prepared as described in *Supporting material*, together with detailed information on the constructs’ DNA sequences.

### Preparation of DNA templates for cDNA display

For the binary model libraries, genes of T26 constructs were amplified by PCR using the corresponding plasmids as templates, and New Left and cnvK_New Ytag primers listed in Table S3 (Fig. S3A). Amplified PCR products were column-purified (QIAquick PCR Purification Kit, QIAGEN, Germany) and their concentration was evaluated by NanoDrop (NanoDrop, USA). Purified DNA was used for *in vitro* transcription.

For the preparation of random libraries, LibQ and Lib4, 61-basepair ssDNA fragments consisting of Gln replaced by degenerate NNK codon, and amino acids at positions -1, +1, +2, +3 from Gln replaced by degenerate NNK codon respectively, were custom ordered from Integrated DNA Technologies, Singapore. The scheme of the library preparation is given in Fig. S3B. dsDNA fragments were generated in the reaction with Klenow fragment (Takara-Bio, Japan) and a single primer, number 16 (Table S3). dsDNA fragments were inserted into PCR-amplified (primers 17-18 in Table S3) and purified pRSET vector fragment by Gibson Assembly (NEB, USA) to add sequence parts necessary for cDNA display generation to the DNA library. Assembled products were PCR amplified using New Left and cnvK_New Ytag primers, column-purified and used for *in vitro* transcription to obtain LibQ and Lib4 mRNA libraries.

### Preparation of cDNA display

mRNA pools were prepared by *in vitro* transcription using the RiboMAX Large Scale RNA Production System-T7 (Promega, USA), and prepared DNA templates. For binary model libraries, DNA containing the original T26 sequence and non-substrate sequence (T26A) was mixed in designated molar ratios and used as a template for the synthesis of the mRNA library. For random libraries, prepared dsDNA corresponding to LibQ and Lib4 was used as the template. The reaction mixtures were incubated at 37°C for 2h. This was followed by purification of the synthesized mRNA, including the on-column DNA digestion, with the NucleoSpin RNA kit (Takara-Bio, Japan). The purity and concentration of RNA were checked by NanoDrop and Urea PAGE electrophoresis. Twenty pmol of the synthesized mRNA libraries were hybridized and photo-crosslinked to the cnv-K linker (Fig. S4A), as done before.^(22; 11)^ Six μL of the obtained reaction mixture was used as a template for *in vitro* translation using the reconstituted *E. coli* -based cell-free protein synthesis system (PURE*frex*, GeneFrontier, Japan) in a 25-μL scale. The reaction mixture was incubated at 37°C for 30 min, followed by the addition of EDTA (20 mM) and incubation at 37°C for 5 min to release the ribosomes. In the following, the reactions were centrifuged, and supernatants containing mRNA display complexes (mRNA-linker-protein complex) were collected for purification. Purification was carried out by immobilization of mRNA display molecules to streptavidin-coated magnetic microbeads (Streptavidin MyOne C1, Thermo Fisher, USA) via biotin of the puromycin linker. After immobilization, the beads were washed with binding buffer (10 mM Tris-HCl, pH 8.0, 1mM EDTA, 1M NaCl, 0.1% Tween 20) and 1X ReverTra Ace buffer. Reverse transcription reaction was then carried out to convert mRNA to cDNA using the ReverTra Ace (Toyobo, Japan) reverse transcriptase at 42°C for 30 min on a rotator. Beads were then washed with selection buffer (50 mM Tris-HCl, pH 7.4, 0.5M NaCl, 1mM EDTA, 0.05% Tween 20), and treated with RNase T1 for 15 min at 37°C on a rotator to release formed cDNA display (mRNA-cDNA-linker-protein complex) by digestion of the RNase T1 recognition site included in the linker structure. Supernatant after RNase T1 treatment containing the cDNA display molecules was used for subsequent selection.

### Selection of transglutaminase substrate peptides

The selection was performed on a 100-μL scale, using 30 μL of cDNA display solution per reaction. In addition, each reaction mixture contained 32 mM pentylamine-biotin (Thermo Fisher, USA) as acyl acceptor substrate, 10 mM Tris-HCl, pH 8.0, 15 mM CaCl_2_, 7.5 mM DTT and 0.01 mg/mL recombinant mouse TG2. The reaction was incubated at 37°C for 90 min, followed by ultrafiltration with a 3 kDa cut-off membrane to remove the unreacted pentylamine-biotin. In the following, formed covalent complexes between cDNA display molecules and pentylamine-biotin were pulled from the mixture by immobilization to streptavidin-coated magnetic microbeads (20 μL; Streptavidin MyOne C1, Thermo Fisher, USA). After immobilization, the beads were separated from the supernatant, washed three times with each, 1 mL of the selection buffer v4 (50mM Tris-HCl, pH 7.4, 0.5M NaCl, 1mM EDTA, 0.7% Tween 20) and 100 μL of TE buffer, followed by resuspension in TE buffer.

### Detection of enriched DNA in model libraries

Original cDNA library, beads suspension (enriched library) and supernatant fractions (leftover library) were used for PCR amplification of DNA using the primers Nested_Fw1 and Nested_Rv1 (Table S3, Fig. S5). At the Q2A mutation site, T26A DNA contains a unique restriction site for *Sph* I, which T26 does not have. This property was used to distinguish between PCR-amplified T26 and T26A DNA. Briefly, 5 μL of PCR product was subjected to restriction digestion with *Sph* I and analyzed by agarose gel electrophoresis. The band pattern was compared with that of digested T26 and T26A original DNA to identify the enriched DNA.

### Analysis of the enriched DNA from LibQ and Lib4 libraries

Original cDNA library, enriched library and leftover library were used for PCR amplification using the primers Nested_Fw1 and Nested_Rv1 (Table S3, Fig. S5) to check for the presence of expected DNA band in each of the fractions. For the enriched library, a portion of the PCR mixture after the first nested PCR was used as a template for the second nested PCR with Nested_Fw2 and Nested_Rv2 primers (Table S3, Fig. S5). Amplified DNA was analyzed by electrophoresis. DNA amplified from the original and enriched library was purified and prepared for NGS analysis.

To prepare the original and enriched libraries for sequencing, corresponding DNA was PCR amplified with the following set of primers: NGS prep (T26)_Fw and Nested_Rv1 for Lib4 before and after selection, NGS prep (T26, randomQ, after) and Nested_Rv1 for LibQ after selection, and NGS prep (T26, randomQ, before) and Nested_Rv1 for LibQ before the selection. Since we analyzed the original and enriched LibQ DNA as a single sequencing sample, this PCR amplification step also introduced specific 10 bp tags upstream of the T26 gene to distinguish the sequence reads belonging to the original library from the ones belonging to the enriched library. After the PCR amplification, DNA was column-purified. Original and enriched LibQ DNA was mixed in a 1:1 molar ratio to make one sample for analysis, while original and enriched Lib4 DNA were analyzed as separate two samples. The samples were further prepared and analyzed by pair-end next-generation sequencing (Illumina, NextSeq550) carried out at the Center for Gene Research of Nagoya University.

### Processing of the NGS data

At first, data quality was assessed by Seqkit^(25)^. Quality scores, Q20(%) and Q30(%), were between 93 and 95%, which indicates a low probability of incorrect base identification in our data sets. Trimmomatic^(3)^ was used to remove contaminating adapter sequences. Then, Seqkit was applied for quality filtering with Q set to 25. After quality filtering, 82-83% of total sequences passed the evaluation and were further used. Fastq files were then converted to fasta format, and only forward sequence reads were retained. This is because all of the mutation sites are within the 81bp forward read of the total 160-170bp long DNA. Reads from the sample containing the tagged sequences from LibQ were processed to separate the reads corresponding to the library before and after selection. The statistics of sequence files were monitored after each processing step to ensure sufficient data are retained for the next step. Finally, all bases upstream of the peptide sequence were removed from the reads and the remaining DNA sequence was converted into the amino acid sequence and analyzed by Biopython^(4)^. For Lib4, the final filtering was applied to exclude sequences which do not contain Gln at the second position, since those sequences are not planed according to the library design but could accidentally be present as a result of chemical synthesis of the DNA library and subsequent carryover. The rank list of top 100 enriched peptides further includes filtering of the sequences that show less than 100 reads after the selection, as these were not considered significant. The ranking was made based on the value of the enrichment factor (from highest to lowest).

## Acknowledgment

This work was supported by Grant-in-Aid for Early-Career scientists from Japan Society for the Promotion of Science awarded to J.D. (18K14387). The authors are thankful to Dr. Takashi Kanamori from Gene Frontier, Japan for providing PURE*frex* components, and to Dr. Bo Zhu from Tokyo Institute of Technology for valuable discussions.

