## Supplementary information for "cDNA display coupled with next-generation sequencing for rapid activity-based screening: Comprehensive analysis of transglutaminase substrate preference"

*To whom correspondence should be addressed:

Running title: Screening of transglutaminase substrate preference by cDNA display

### Methods

*Preparation of T26 DNA constructs*

The gene of the T26-GST construct was sub-cloned from pET24d-T26-GST(QN) plasmid to obtain pRSET-T26-GST by the In-fusion cloning kit (Takara-Bio, Japan). pET24d-T26-GST(QN) contains a specially designed GST tag with all Gln residues replaced by Asn to prevent it from becoming a TG substrate. Since we only used this modified GST tag, in the following, the description '(QN)' behind GST will be omitted for simplicity. pRSET vector backbone and T26-GST gene insert were PCR-amplified with primers 1-4 listed in Table S3 using pRSET-P450 and pET24d-T26-GST as templates respectively. Amplified products were purified and fused to obtain a pRSET-T26-GST plasmid. As a negative control, pRSET-T26N-GST with Q2N mutation was obtained by inverse PCR using pRSET-T26-GST as a template and mutagenic primers 5-6 listed in Table S3, followed by self-circularization with In-fusion cloning kit. In the following, another inverse PCR step with pRSET-T26N-GST as a template and mutagenic primers 7-8 listed in Table S3 was performed to introduce restriction site for *Xho*I in the form of silent mutation in the sequence of the GST tag. The final plasmid was obtained by subsequent self-circularization with In-fusion cloning kit.

Untagged T26 construct was prepared by inverse PCR with primers 9-10 listed in Table S3, using pRSET-T26-GST plasmid as a template. This was followed by ligation using Mighty Mix (Takara-Bio, Japan).

Non-substrate peptide, the untagged T26A construct with Q2A mutation, was obtained by inverse PCR with mutagenic primers 11-12 listed in Table S3 and pRSET-T26 (long linker) as a template. In-fusion cloning was used to self-circularize the obtained DNA. Introduced Gln to Ala mutation resulted in the introduction of the *Sph*I restriction site.

The outline of all used constructs is given in Figure S1.

**Results**

*Display of T26 sequence genetically fused with GST*

Previous studies on the characterization of isolated TG2 Gln probes used C-terminal GST-tag for soluble expression of peptides in *E.coli*. We have initially followed this methodology and displayed T26 as a protein fusion with the C-terminal GST-tag.

Genes of T26-GST and T26N-GST were amplified by PCR using the corresponding plasmids as templates and a combination of New Left and New Ytag primers listed in Table S3. After PCR amplification, the products were column-purified and used as templates for synthesis of mRNA by *in vitro* transcription. mRNA was hybridized and enzymatically ligated to SBP puromycin linker (Fig. S4B).

Selection of T26-GST from the 1:1 binary model library made by mixing T26-GST and T26N-GST cDNA display solutions in a 1:1 ratio was a success, with only T26-GST DNA detected in the enriched pool (Fig. S2A). In the following, we made 1:10 and 1:100 model libraries by mixing the T26-GST and T26N-GST DNA in designated ratios before *in vitro* transcription to mimic the real library design. However, the enrichment was unsuccessful (Fig. S2B).

**Figures and tables**

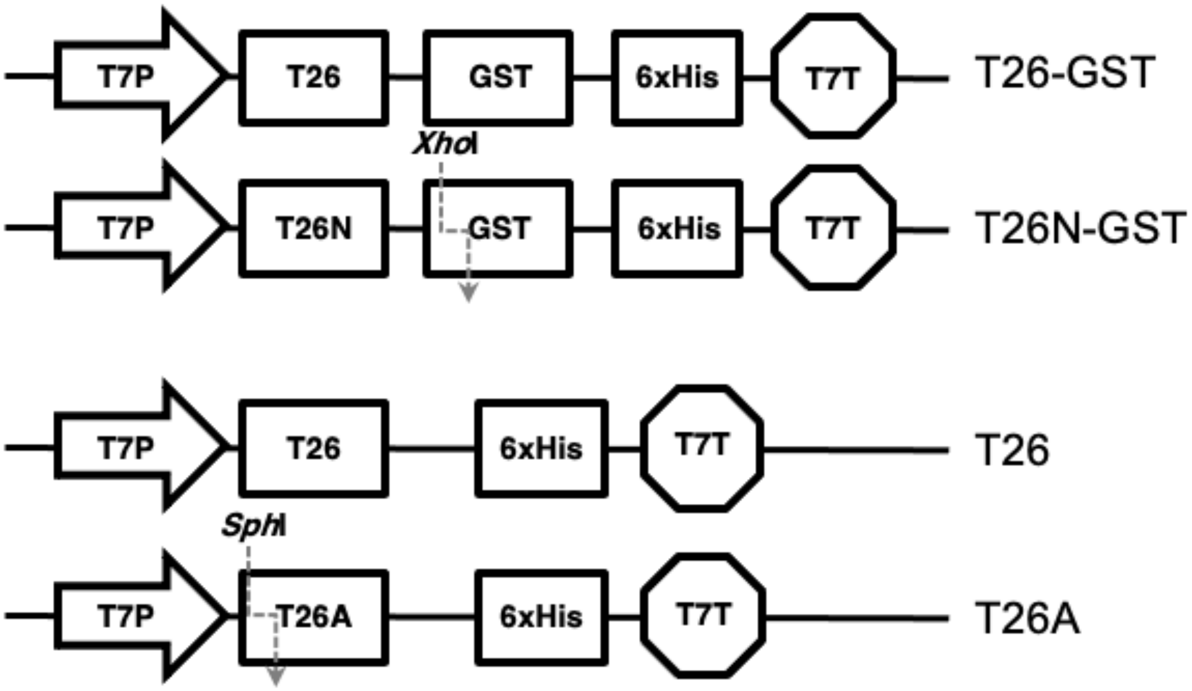


**Figure S1.** DNA constructs used in this study. Vector backbone (pRSET) was the same for all constructs and thus is omitted.

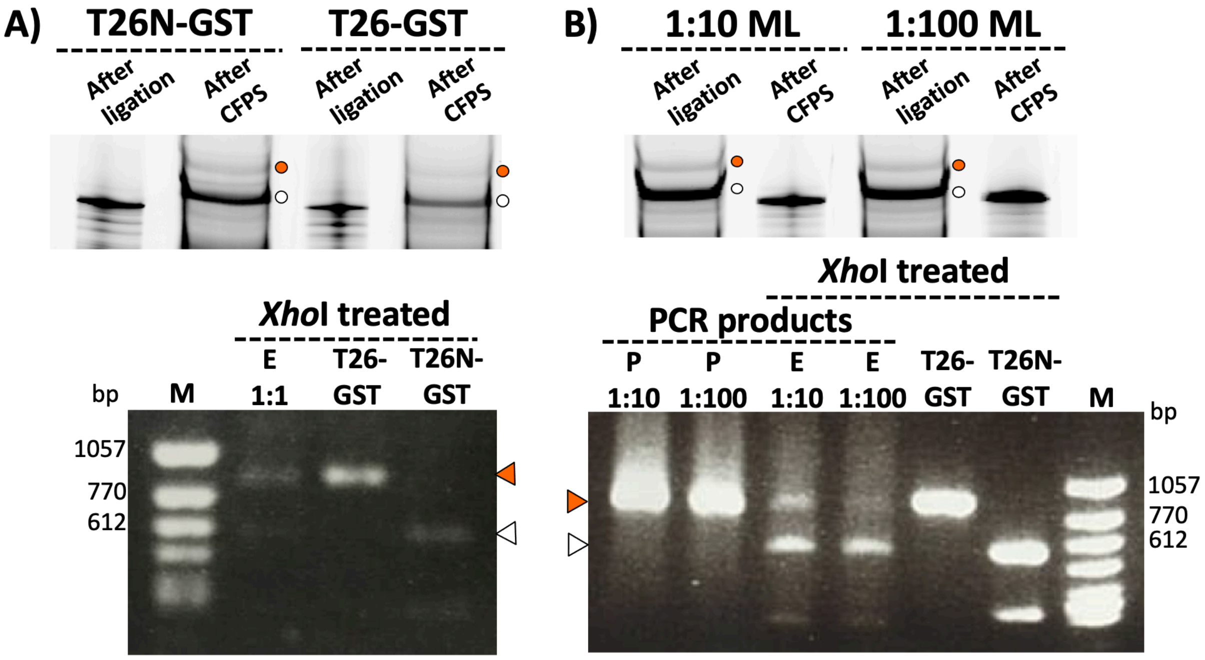


**Figure S2.** Upper panels: Urea SDS-PAGE gel of the ligation products and mRNA display complexes visualized using fluorescence imager under the FITC filter. Orange circles indicate the position of mRNA display complexes and white circles indicate the position of ligation products (mRNA-linker complexes). Lower panels: Electrophoresis result of the DNA analysis after selection, PCR amplification (P) and *Xho*I treatment (E). Orange triangles indicate the position of the T26-GST DNA band, and white triangles indicate the position of the T26N-GST DNA band. T26-GST and T26N-GST are lanes with the corresponding DNA after *Xho*I treatment used as a reference for the evaluation of enriched DNA composition. (A) Results from 1:1 model library. (B) Results from 1:10 and 1:100 model library.

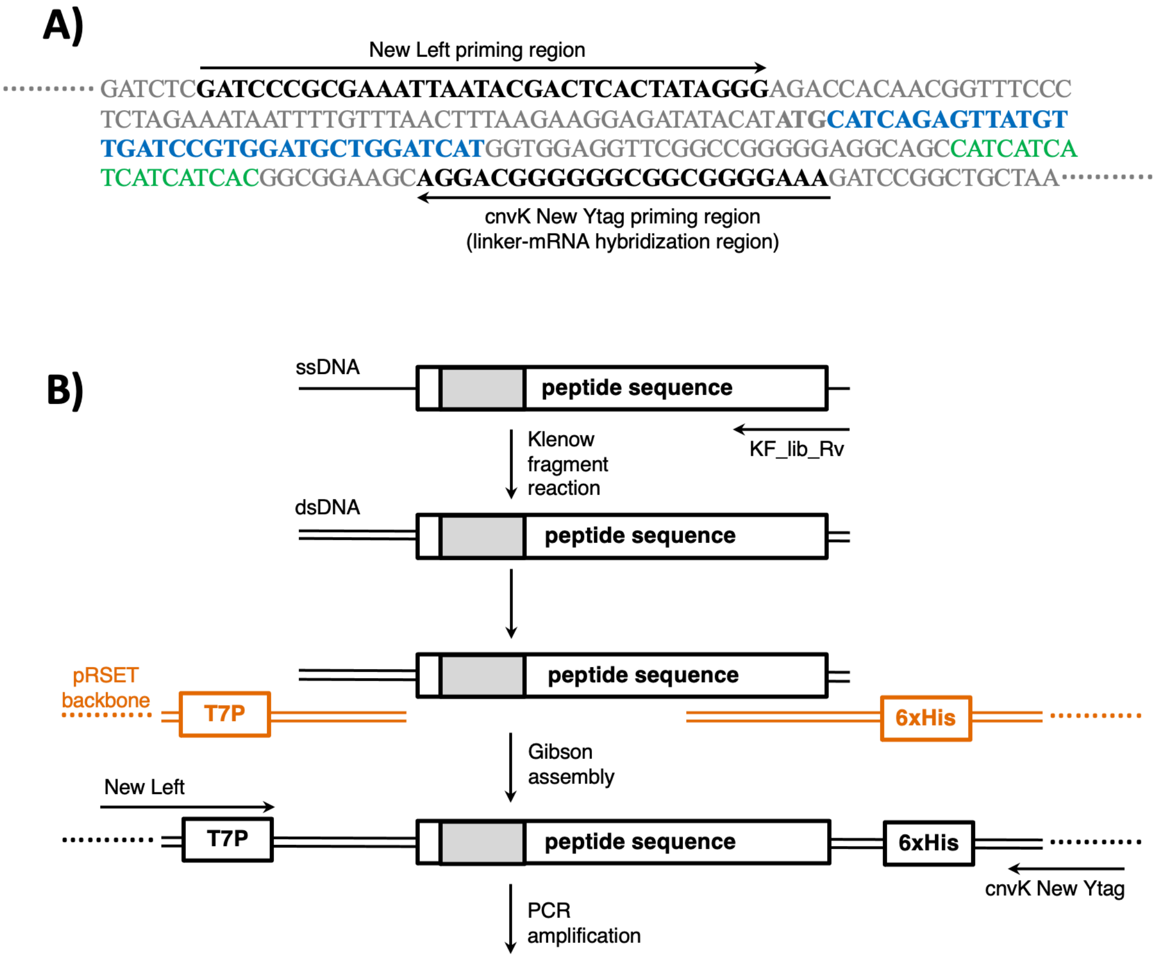


**Figure S3.** (A) Preparation of the dsDNA template for cDNA display by PCR with New Left and cnvK_New Ytag primers; (B) Preparation of the dsDNA template of the random libraries, LibQ and Lib4.

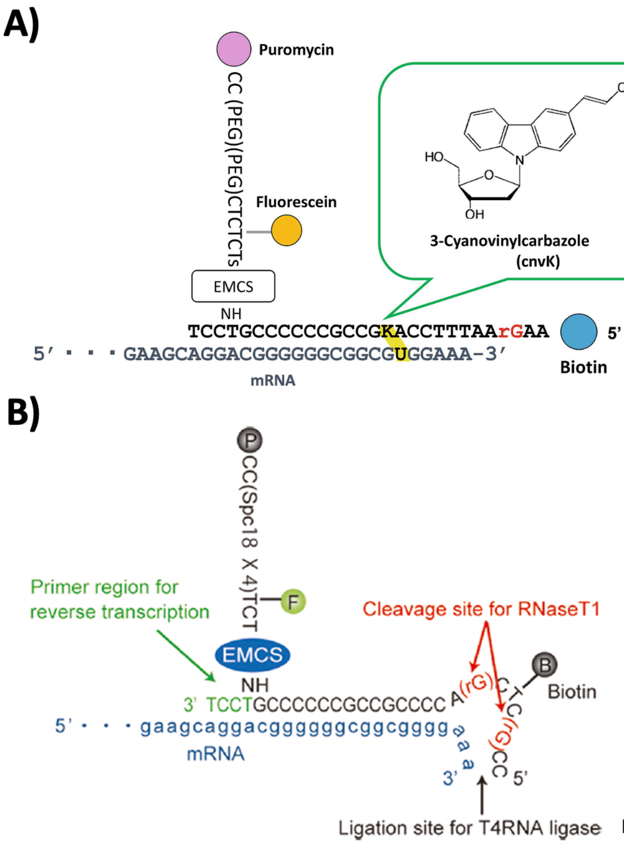


Nemoto *et al*. (2018) Antibody Engineering: Methods and Protocols., Methods Mol. Biol., Vol.1827, New York, NY: Springer New York. p 269-285.

Mochizuki *et al*. (2011) ACS Combinatorial Science 13:478-485.

**Figure S4.** Structures of the puromycin linkers, cnvK linker (A) and SBP linker (B), used in this study.

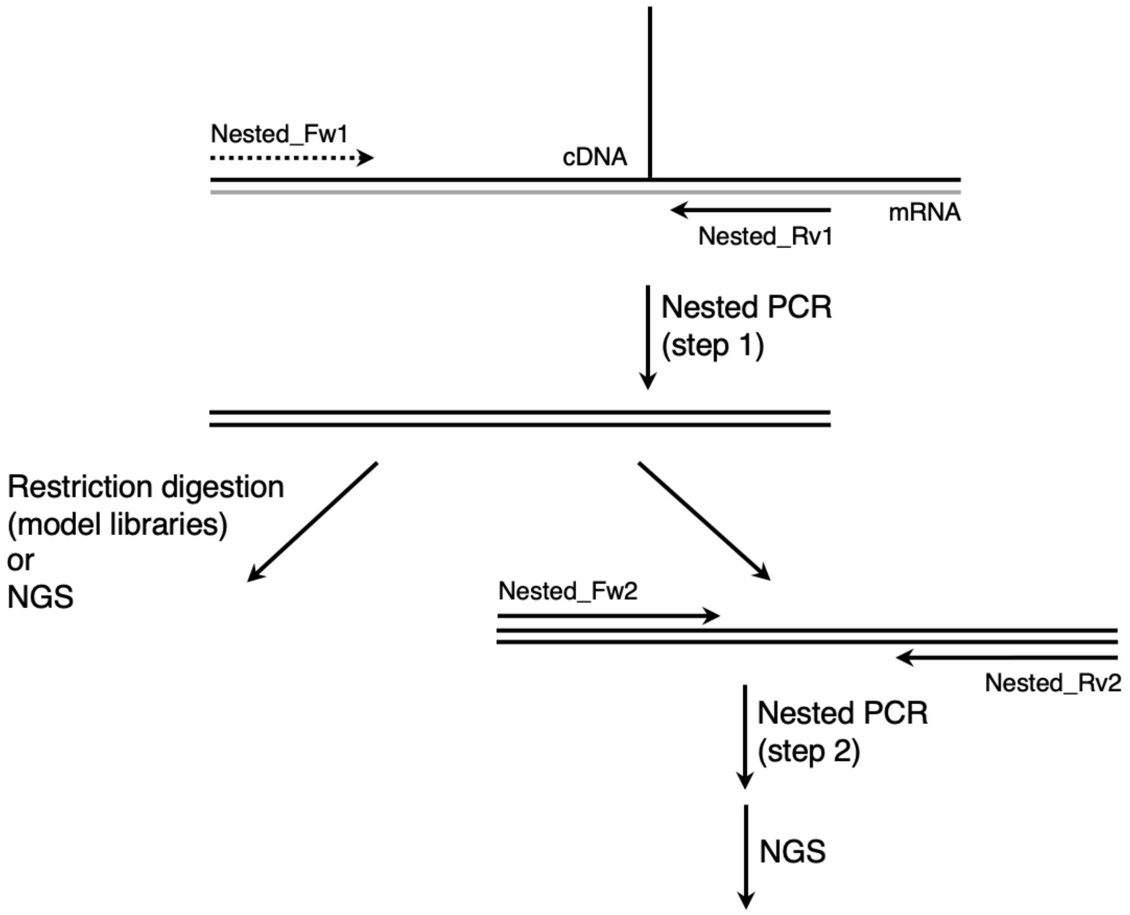


**Figure S5.** Post-selection preparation of the enriched DNA for analysis. The first step involves PCR, after which DNA is either used for analysis by restriction digestion (model libraries) and NGS (original LibQ and Lib4, before the selection), or for the second PCR (enriched complexes from LibQ and Lib4, after the selection) followed by NGS.

**Table S1**. DNA sequences of the cDNA display constructs used in this study. T7P sequence is bolded, peptide sequence is colored red, and 6xHis sequence blue. Randomized sites are italicized.

| Name | Sequence (5'-3') |
| --- | --- |
| T26 | GATCCCGCGAAAT**TAATACGACTCACTATAGG**GAGACCACAACGGTTTCCCTCTAGAAATAATTTTGTTTAACTTTAAGAAGGAGATATACATATG**CATCAGAGTTATGTTGATCCGTGGATGCTGGATCAT**GGTGGAGGTTCGGCCGGGGGAGGCAGCCATCATCATCATCATCACGGCGGAAGCAGGACGGGGGGCGGCGTGGAAA |
| T26A | GATCCCGCGAAAT**TAATACGACTCACTATAGG**GAGACCACAACGGTTTCCCTCTAGAAATAATTTTGTTTAACTTTAAGAAGGAGATATACATATG**CATGCGAGTTATGTTGATCCGTGGATGCTGGATCAT**GGTGGAGGTTCGGCCGGGGGAGGCAGCCATCATCATCATCATCACGGCGGAAGCAGGACGGGGGGCGGCGTGGAAA |
| LibQ | GATCCCGCGAAAT**TAATACGACTCACTATAGG**GAGACCACAACGGTTTCCCTCTAGAAATAATTTTGTTTAACTTTAAGAAGGAGATATACATATG**CAT*NNK*AGTTATGTTGATCCGTGGATGCTGGATCAT**GGTGGAGGTTCGGCCGGGGGAGGCAGCCATCATCATCATCATCACGGCGGAAGCAGGACGGGGGGCGGCGTGGAAA |
| Lib4 | GATCCCGCGAAAT**TAATACGACTCACTATAGG**GAGACCACAACGGTTTCCCTCTAGAAATAATTTTGTTTAACTTTAAGAAGGAGATATACATATG***NNK*CAG*NNKNNKNNK*GATCCGTGGATGCTGGATCAT**GGTGGAGGTTCGGCCGGGGGAGGCAGCCATCATCATCATCATCACGGCGGAAGCAGGACGGGGGGCGGCGTGGAAA |

**Table S2.** Rank list showing the top 100 peptide sequences enriched from Lib4.

| Ranking | Peptide sequence | Enrichment factor | Sequence count |
|---|---|---|---|
| 1 | QQVCIDP | 7.12 | 112 |
| 2 | QQYVVDP | 6.62 | 125 |
| 3 | QQFVVDP | 6.17 | 110 |
| 4 | QQFRVDP | 5.72 | 150 |
| 5 | QQVWVDP | 5.35 | 129 |
| 6 | QQYRVDP | 5.31 | 156 |
| 7 | WQTWVDP | 5.22 | 115 |
| 8 | PQVWFDP | 5.10 | 107 |
| 9 | AQWYVDP | 5.10 | 139 |
| 10 | CQFAMDP | 5.06 | 122 |
| 11 | HQFAVDP | 4.96 | 130 |
| 12 | MQYAVDP | 4.94 | 114 |
| 13 | CQTWIDP | 4.92 | 129 |
| 14 | SQCWIMV | 4.90 | 257 |
| 15 | QQYLVDP | 4.90 | 149 |
| 16 | QQYGLDP | 4.87 | 138 |
| 17 | MQFAVDP | 4.86 | 107 |
| 18 | WQHVVDP | 4.48 | 155 |
| 19 | CQKCVDP | 4.47 | 122 |
| 20 | QQIVVDP | 4.46 | 145 |
| 21 | AQCAIDP | 4.42 | 116 |
| 22 | QQVICDP | 4.33 | 100 |
| 23 | QQCRVDP | 4.29 | 171 |
| 24 | QQVILDP | 4.22 | 115 |
| 25 | CQYRIDP | 4.21 | 212 |
| 26 | QQSSIDP | 4.20 | 119 |
| 27 | CQCAIDP | 4.17 | 127 |
| 28 | CQTWVDP | 4.17 | 162 |
| 29 | CQYWVDP | 4.17 | 162 |
| 30 | QQIGVDP | 4.14 | 100 |
| 31 | WQCRIDP | 4.14 | 126 |
| 32 | YQFEVDP | 4.10 | 116 |
| 33 | QQVIVDP | 4.07 | 175 |
| 34 | AQYWVDP | 4.03 | 203 |
| 35 | SQQCVDP | 4.01 | 181 |
| 36 | YQYRCDP | 4.01 | 101 |
| 37 | YQCWIMV | 3.99 | 134 |
| 38 | WQCHVDP | 3.97 | 100 |
| 39 | WQSIIDP | 3.95 | 112 |
| 40 | CQYALDP | 3.95 | 174 |
| 41 | QQWVLDH | 3.95 | 116 |
| 42 | GQKYVDP | 3.94 | 161 |
| 43 | GQCWIMV | 3.93 | 136 |
| 44 | TQCAVDP | 3.93 | 136 |
| 45 | WQYRIDP | 3.93 | 140 |

| | | | |
|---|---|---|---|
| 46 | YQITVDP | 3.91 | 123 |
| 47 | AQHRVDP | 3.91 | 127 |
| 48 | AQEWVDP | 3.88 | 122 |
| 49 | GQQVIDP | 3.88 | 122 |
| 50 | WQCEVDP | 3.87 | 138 |
| 51 | QQVYVDP | 3.84 | 129 |
| 52 | QQYRTDP | 3.81 | 100 |
| 53 | QQSWVDP | 3.81 | 128 |
| 54 | VQQVIDP | 3.81 | 136 |
| 55 | MQVWIDP | 3.79 | 139 |
| 56 | WQFTVDP | 3.78 | 123 |
| 57 | WQCVIDP | 3.78 | 115 |
| 58 | QQVRCDP | 3.78 | 107 |
| 59 | AQYVVDP | 3.76 | 308 |
| 60 | QQIVLDH | 3.75 | 189 |
| 61 | QQVRIDP | 3.75 | 114 |
| 62 | WQTVCDP | 3.75 | 110 |
| 63 | QQSRVDP | 3.73 | 141 |
| 64 | CQVWVDP | 3.71 | 370 |
| 65 | CQYTIDP | 3.71 | 109 |
| 66 | MQFGIDP | 3.71 | 109 |
| 67 | GQLCWIM | 3.70 | 101 |
| 68 | YQYCVDP | 3.69 | 120 |
| 69 | CQYAMDP | 3.68 | 112 |
| 70 | QQYVSDP | 3.68 | 108 |
| 71 | YQLCVDP | 3.66 | 234 |
| 72 | CQQSVDP | 3.65 | 161 |
| 73 | QQVRVDP | 3.65 | 264 |
| 74 | GQFTVDP | 3.63 | 335 |
| 75 | YQVHIDP | 3.63 | 118 |
| 76 | WQVQLDP | 3.62 | 133 |
| 77 | SQQYVDP | 3.62 | 148 |
| 78 | AQFAVDP | 3.60 | 102 |
| 79 | QQLWIDP | 3.60 | 102 |
| 80 | CQIVCDP | 3.60 | 136 |
| 81 | QQIGLDP | 3.59 | 113 |
| 82 | TQYAVDP | 3.59 | 113 |
| 83 | GQYQVDP | 3.59 | 143 |
| 84 | CQCVIDP | 3.59 | 267 |
| 85 | CQTRIDP | 3.57 | 161 |
| 86 | GQHVIDP | 3.57 | 187 |
| 87 | GQCMIDP | 3.56 | 127 |
| 88 | MQFVVDP | 3.55 | 220 |
| 89 | CQLGIDP | 3.54 | 334 |
| 90 | VQYCIDP | 3.53 | 215 |
| 91 | CQMCIDP | 3.53 | 111 |
| 92 | CQIRIDP | 3.52 | 166 |
| 93 | VQCAIDP | 3.52 | 166 |

| | | | |
|---|---|---|---|
| 94 | CQYHVDP | 3.51 | 129 |
| 95 | QQYVLDP | 3.51 | 199 |
| 96 | HQYAVDP | 3.51 | 103 |
| 97 | WQIRVDP | 3.50 | 220 |
| 98 | NQFAVDP | 3.50 | 121 |
| 99 | YQAKVDP | 3.49 | 128 |
| 100 | CQWSIDP | 3.48 | 212 |

* N-terminal Met is present in all peptides and represents a translated start codon which remained uncleaved due to the absence of methionine aminopeptidase in the PURE system. It is omitted from the peptide sequences in the list since it does not represent preference of TG2.

**Table S3.** List of oligonucleotide primers used in this study.

| Number | Name | Sequence (5'-3') | Use |
|---|---|---|---|
| 1 | In-fusion gene_Fw | CATCAGAGTTATGTTGATCCGTGGATG | Cloning of T26-GST gene into the pRSET |
| 2 | In-fusion gene_Rv | TTTTGGAGGATGGTCGCCACCAC | |
| 3 | In-fusion vector_Fw | GACCATCCTCCAAAAGGGGGAGGCAGCCATCATCAT | |
| 4 | In-fusion vector_Rv | AACATAACTCTGATGCATATGTATATCTCCTTCTTAAAGTTAAACAAA | |
| 5 | In-fusion T26QN_Fw | GATATACATATGCATAACAGTTATGTTGATCCGTGG | Preparation of pRSET-T26N -GST |
| 6 | In-fusion T26QN_Rv | ATGCATATGTATATCTCCTTCTTAAAGTT | |
| 7 | Leu_sil_cut_site_Fw | ATTGGGTCTCGAGTTTCCCAATCTTCCTTATT | Introduction of *Xho*I restriction site into pRSET-T26N-GST |
| 8 | Leu_sil_cut_site_Rv | AACTCGAGACCCAATTCAAACTTTTTGTTTCG | |
| 9 | T26 only_Fw | GGGGGAGGCAGCCATCATCATCATC | Preparation of pRSET-T26 |
| 10 | T26 tag (vector)_Rv | GGCCGAACCTCCACCATGATCCAG | |
| 11 | T26 (long linker)_QtoA_Fw | GCGAGTTATGTTGATCCGTGGATGCTG | Preparation of pRSET-T26A |
| 12 | T26 (long linker)_QtoA_Rv | ATCAACATAACTCGCATGCATATGTATATCTCC | |
| 13 | New Left | GATCCCGCGAAATTAATACGACTCACTATAGGG | Preparation of DNA template for cDNA display |
| 14 | New Ytag | TTTCCCCGCCGCCCCCCGTCCT | |
| 15 | cnvK_New Ytag | TTTCCACGCCGCCCCCCGTCCT | |
| 16 | KF_lib_Rv | CCATGATCCAGCA | Klenow fragment reaction |
| 17 | GA_vec_lib_Fw | GTGGATGCTGGATCATGGTGGA | pRSET preparation for Gibson assembly to dsDNA library |
| 18 | GA_vec_lib_Rv | TGTATATCTCCTTCTTAAAGTTAAACAAAATTAT | |
| 19 | Nested_Fw1 | GGGAGACCACAACGGTTTCC | Amplification of DNA after selection |
| 20 | Nested_Rv1 | TTTCCCCGCCGCCCCC | |
| 21 | Nested_Fw2 | GGGAGACCACAACGGTTTCCCTCTAGAAAT | |
| 22 | Nested_Rv2 | TTTCCCCGCCGCCCCCCGTC | |
| 23 | NGS prep (T26)_Fw | CCCTCTAGAAATAATTTTGTTTAACTTTAAG | Preparation of libraries for NGS |
| 24 | NGS prep (T26, randomQ, after) | CTGACAAAAACCCTCTAGAAATAATTTTGTTTAACTTTAAG | |
| 25 | NGS prep (T26, randomQ, before) | CTGACTTTTTCCCTCTAGAAATAATTTTGTTTAACTTTAAG | |
